# Multi-Dimensional Spatiotemporal Attention Neural Network for Next Generation Sequencing Basecalling

**DOI:** 10.1101/2025.11.06.686985

**Authors:** Kuankuan Peng, Wei Chen, Tianran Yao, Huihua Xia, Guoli Fu, Gailing Li, Yuanye Bao, Erkai Liu, Luyang Zhao, Gufeng Wang

## Abstract

Next-generation sequencing (NGS) remains the most used sequencing technique in the field of genomics. Traditional basecall methods face significant challenges in decoding high density sequencing data due to inherent noise in biochemical reactions and limitations of instruments. Here, we present a multi-dimensional deep learning neural network based on spatiotemporal attention mechanism named AICall. The network skips computationally heavy but less effective steps of peak finding and brightness extraction/correction, and directly basecalls from the time sequence of multi-dimensional image stacks obtained in real time. By introducing attention mechanism, it effectively extracts spatial and time-related key information including spatial crosstalk, spectral crosstalk, phasing, base-quenching, and intensity decay, and significantly improves basecall accuracy. We demonstrate that AICall achieves an average error rate less than 0.01% and provides more reliable sequencing results for downstream analysis.

---

Next generation sequencing has had a profound impact on our daily life. Compared to Sanger sequencing, its overall error rate is relatively high, on the 0.1% level. This is unfavorable for many applications that require the detection of minor base mutations. There are recent methods using *a posteriori* information to re-calibrate the Phred Q-values of measured bases; high quality calls are selected and assigned a high Q score such as Q40 or even higher. However, erroneous basecalls are NOT corrected and the overall error rate remain on the 0.1% level.

Traditional NGS basecall algorithm, taking 4-colored sequencing as an example, includes peak finding, image registration, intensity extraction and correction, and base recognition of the 4-colored images. In particular, basecall accuracy solely relies on thus is extremely sensitive to the fluorescence intensity of the clusters where the sequencing-by-synthesis reaction takes place. However, due to biochemical reaction inhomogeneity and instrument limit, the fluorescence intensity of the clusters is affected by many interference factors such as template copy number variation between clusters, templates’ reactions out of sync with in a cluster (phasing/pre-phasing), spectral crosstalk, spatial crosstalk, intensity decay over cycles, G base-quenching, etc. In addition, Signal is further subjected to distortion during the intensity extraction step because of (1) systematic reasons: finite size of camera sensing pixels and imperfect alignment of the 4-color images due to systematic optical aberrations (chromatic and Seidel); and (2) random causes: e.g., local temperature field fluctuation causing random image deformation.

Current base-calling algorithms can be classified into two categories. The first type models measured signal as the product of the “true” signal and the influences, often formulated as matrices that apply on the “true” signal vector. Through regression or fitting of the experimental data, the interference matrices can be solved, leading to the recovery of the “true” signal. Most of the NGS manufacturers adopt this approach in whole or in part, as it effectively corrects the influence from phasing/pre-phasing and spectral crosstalk. However, the spatial crosstalk is challenging to model when convoluted with optical aberrations, which is usually not regular in the large field of view; random image deformation worsens the situation. While manufacturers are pushing the limit of the density of current sequencing flow cells, the above two problems combined appear to be one of the most important bottlenecks of current NGS sequencing technology.

The second type is based on machine learning, which relies on supervised training to learn the complex bias patterns during basecall. Most of these methods use extracted intensity for training and basecalling, meaning errors are introduced from the beginning of the analysis. Recently, deep learning methods have been explored for sequencing basecalling. A hierarchical DNN method used CNN to locate the cluster centers and extract the signals, which are then fed into a DNN for base classification. Another method called Baseformer uses CNN to locate the centers of clusters and encodes the cluster with its surrounding pixels as input to a Transformer model for cluster classification. Both methods leverage information from neighboring pixels of the cluster center and lessens the impact of imperfect image alignment. Yet they process clusters independently, failing to capture the interference among clusters in the intertwined spatial crosstalk network.

## Result

### Neuron network structure

To address the aforementioned challenges, we developed a novel network model for base-calling (AICall, Fig. 1). In contrast to early methods that model only the pixel centroids of clusters, AICall utilizes a stack of fluorescent images as its input (Fig. 1abc). This strategy benefits the model by learning both systematic and random interferences in the multi-dimensional space: xy-, spectral- and temporal dimensions. Three modules were designed to undertake the interference in the multi-dimensional space: Gene Fluorescence Attention (GFA) Module, which includes Gene Fluorescence Spatial Attention (GFSA) and Gene Fluorescence Channel Attention (GFCA) sub-modules (Fig. 1b); Sequence Encoding Module (SEM); and Sequence-Aware Residual Block (SARB) (Fig. 1d).

**Fig. 1.**
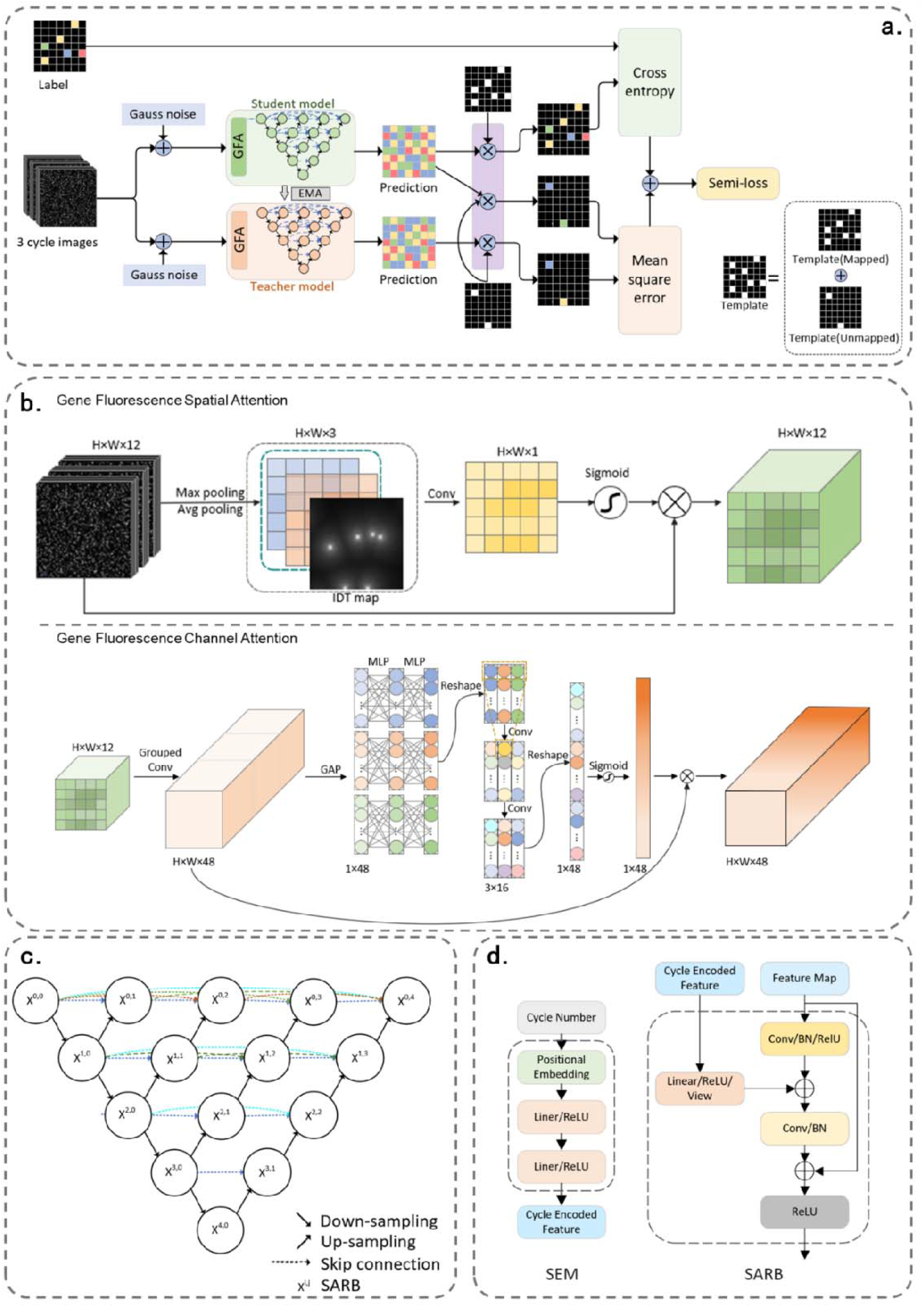
Architecture of deep neural network (DNN) in AICall. (a) DNN semi-supervised training architecture. (b) Gene Fluorescence Attention Module (GFA), encompassing Gene Fluorescence Spatial Attention and Gene Fluorescence Channel Attention. (c) UNet++ architecture. UNet++ is composed of an encoder and a decoder, which are linked by a sequence of nested SARBs. x_i,j_ represents the output of node X_i,j_, where i denote the down-sampling layer along the encoder and j denotes SARB along the skip pathway. (d) Illustration of SEM and SARB.

In the spatial dimensions, we generated inverse distance transformation (IDT) maps based on the cluster location information from the fluorescence images (Fig. 1b upper panel). These maps were then combined with a traditional spatial attention mechanism to construct the GFSA. By enhancing the feature representation of key locations while suppressing irrelevant regions, GFSA effectively improves the model’s feature extraction capability and mitigates the impact from both spatial crosstalk and intensity distortions, which are challenging to model in traditional approaches.

In the spectral (also named as color channel, abbreviated as channel) and local time dimensions, we first employed group convolution, by treating each cycle as an individual group to extract intra-cycle feature maps (Fig. 1b lower panel). Then, we processed the channel features from three consecutive cycles using three independent Multi-Layer Perceptrons (MLPs), allowing the channel attention mechanism to focus within each cycle. Subsequently, a convolutional layer is introduced to facilitate cross-cycle interactions through the convolution’s local receptive field, capturing feature dependencies and correlations across cycles. This design enables GFCA to differentiate and focus on both intra-cycle and inter-cycle features, addressing interference factors such as spectral crosstalk, phasing/pre-phasing, and G-quenching etc.

In the long scale time dimension, SEM transforms discrete cycle indices into unique high-dimensional feature vectors, which are then fused with the image feature maps in the SARB (Fig. 1d). This mechanism enhances the neural network’s sensitivity to cycle-dependent information by automatically adjusting the feature weights for each cycle. It effectively corrects the influence of biochemical factors such as fluorescence decay, as well as phasing/pre-phasing which also varies over time.

Finally, in the training, precise annotation of data is crucial. Base labels in a read were generated using that called by traditional methods and corrected by mapping against the reference. However, the shortcoming of this approach is that only a part of the reads can be recovered. Those low quality ones are usually discarded, leading to a biased training favoring easy bases while the performance for challenging data is poor. To overcome this problem, we employed a semi-supervised learning framework, which consists of two sub-models: the student model and the teacher model (Fig. 1a). They share the same network architecture but do not share parameters. For labeled bases, the model convergence is guided by computing the cross-entropy loss between the student model’s predictions and the ground truth. For unlabeled bases, consistency is enforced by calculating the mean square error (MSE) between the predictions of the teacher model and the student model.

### Initial evaluation of basecall accuracy

Since original image data for commercial sequencers are not accessible, we collected data from 30 sequencing runs generated on 5 different Salus Pro sequencers. Traditional basecaller SalusCall was used for benchmarking. SalusCall is developed by Salus Medical Technology Inc. Ltd. and has been integrated into the Salus Pro sequencers. This product has been commercially released and scrutinized by users in the market. The data quality is comparable to those from Illumina’s Novaseq 6000 sequencing platform.

Initial evaluation of the basecaller was based on two key metrics of gene sequencing quality: “crude mapping rate” and “crude mismatch rate” against the reference. We conducted 30 sequencing runs of *E coli* (ATCC8739 strain) at different cluster densities. From Fig. 2, it is evident that AICall consistently exhibits better performance in both metrics, with the largest improvement for dense clusters. (A cluster is defined as a dense cluster if there is another cluster has a distance <=2.0 pixel on camera chip. The imaging system parameter: peak fwhm 700 nm at 700 nm wavelength, size of a chip is 246 nm.) The average crude mismatch rate decreased by a factor of ∼2.3, from 0.54% to 0.23% (Tabel 1).

**Fig. 2.**
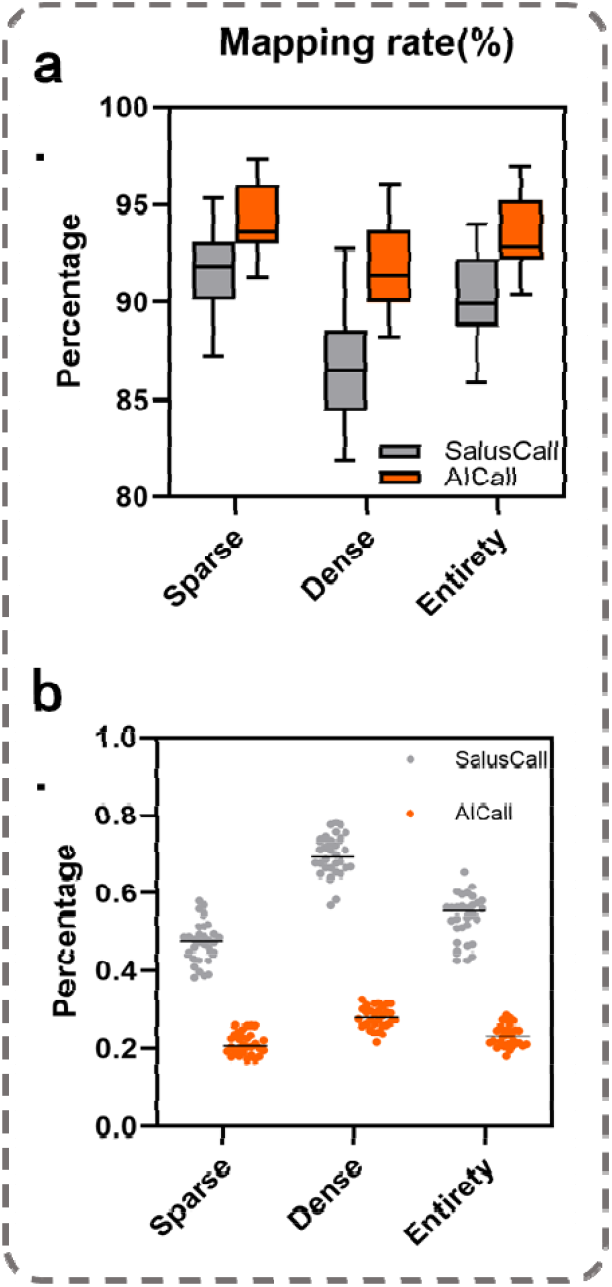
Comparison of the overall performance between AICall and SalusCall on dataset from 30 sequencing runs. (a) Box plots illustrating the distribution of mapping rates for AICall and SalusCall sequencing results, with the center line representing the median, and the box limits indicating the upper and lower quartiles. (b) Scatter plots showing the distribution of mismatch rates for AICall and SalusCall sequencing results, where the center line denotes the median.

Please note that the parameters in Table 1 are crude results for all recovered clusters that contain a portion of low-quality ones. Commercial instrument manufacturers usually remove poor reads before publishing the results. Thus, these parameters are NOT meant to be compared with “mapping rate” and “mismatch rate” for data output by a commercial sequencer. Please see later sections for “mapping rate” and “mismatch rate” of the data output by AICall.

**Table 1.** Average “crude mapping rate” and “crude mismatch rate” of AICall and SalusCall on the test set, including separate statistics for sparse and dense clusters.

| Methods | Crude mapping rate (%) |  |  | Crude mismatch rate (%) |  |  |
| --- | --- | --- | --- | --- | --- | --- |
|  | Sparse | Dense | Average | Sparse | Dense | Average |
| SalusCall | 94.36 | 91.71 | 93.58 | 0.4719 | 0.6948 | 0.5371 |
| AICall | 91.69 | 86.72 | 90.24 | 0.2132 | 0.2813 | 0.2334 |

In achieving above results, we designed a set of metrics and evaluated AICall’s impact on different types of interference, which provided guidance for our model optimizations.

### Spatial crosstalk and intensity distortion caused by imperfect alignment

For a random-cluster sequencing chip, the distance between clusters may be too close that their fluorescence intensities interfere with each other, leading to miscalls. Fig. 3a shows a typical case where an “A” base (red cross) is miscalled as a “T” base (blue cross) because of the strong spatial crosstalk by traditional baseballers. Fig. 3b illustrates how two closely spaced clusters interfere with each other: in any color channel, e.g. “A”, if we plot cluster 1 intensity (y-axis) vs. cluster 2 intensity (x-axis), we will see scattered dots forming 3 lines. The more horizontal line represents cluster 1-X: cluster2-A case (X denotes a non-A base); the more vertical line represents cluster 1-A: cluster 2-X case; the diagonal represents both A case; and the origin spot represents neither A case. The dots are scattered both axially and transversely along the line because of systematic and random interferences.

**Fig. 3.**
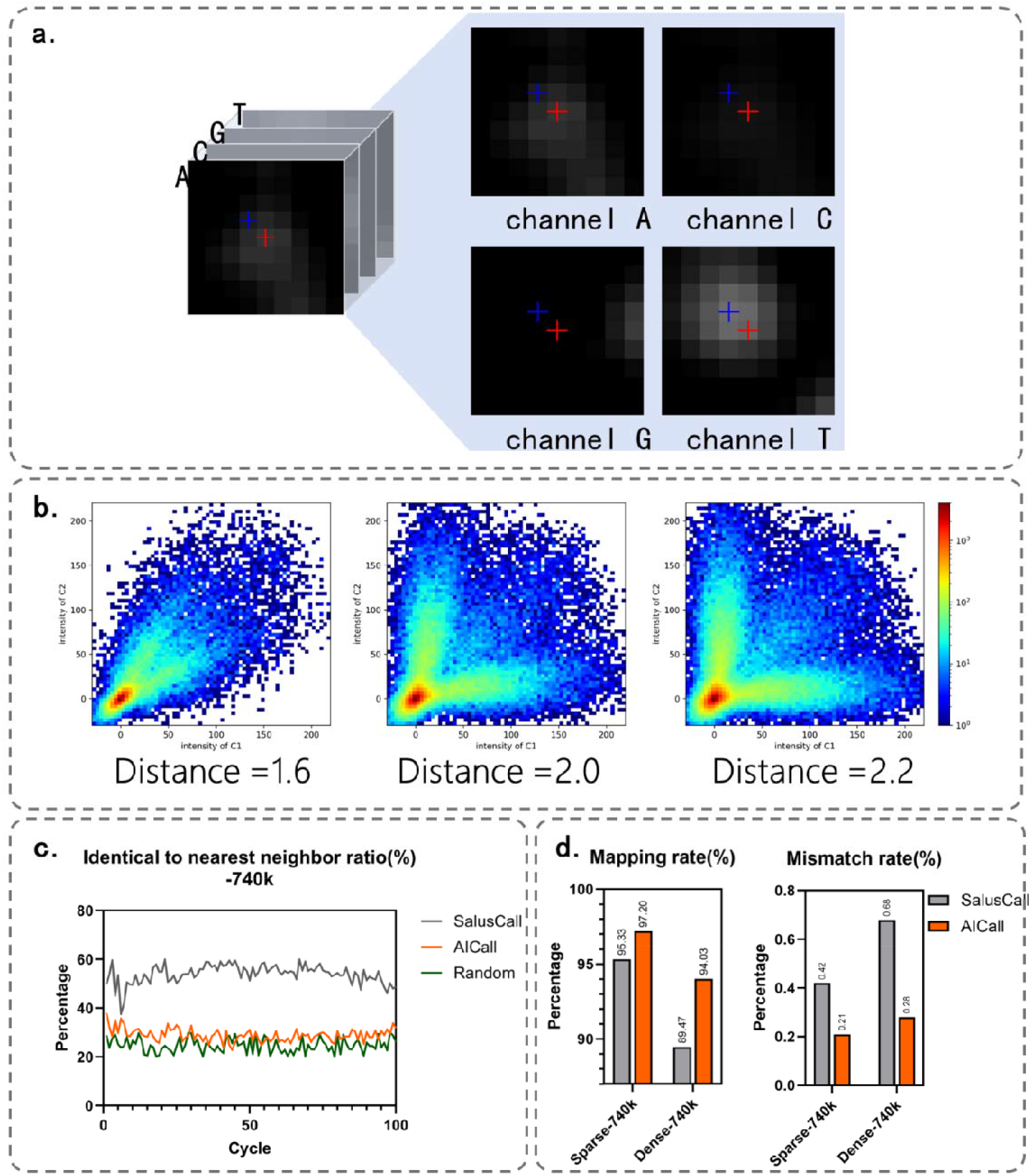
Spatial crosstalk correction. (a) Spatial crosstalk between two clusters. (b) Th relationship between the degree of brightness crosstalk and the distance between two clusters, the x-axis represents the brightness of Cluster 2, while the y-axis represents the brightness of Cluster 1. (c) The proportion of mismatched clusters in AICall and SalusCall sequencing results that ar identical to the base type of their nearest neighbor. The green line represents the proportion of mismatched clusters that have the same base as random clusters. (d) Comparison of AICall and SalusCall sequencing results on sparse clusters and dense clusters.

As the clusters draw near, the horizontal and vertical lines will be closer to the diagonal. Consequently, one cluster is more inclined to interfere with the basecall of the other. As the two clusters are separated by 2.2 camera pixels, there is almost no spatial interference. We arbitrarily selected 2.0 pixels as the threshold to defined “dense cluster” (<=2.0) and “sparse cluster” (>2.0).

We tested samples at different densities (e.g., 620K,740K and 870K per millimeter squared). At a density of 870K/mm^2^, dense clusters account for approximately 30-40% of the total. Fig. 3d shows that both SalusCall and AICall performs worse at high densities. However, AICall out performs SalusCall significantly, by a factor of 2.4 lower in mismatch rate.

To have a more intuitive observation of the impact of spatial crosstalk on basecall and AICall, we plotted the percentage of miscalled dense clusters identical to their nearest neighbor at each of the 100 sequencing cycles (Fig. 3c). It shows that ∼50% of miscalled clusters match their neighbor by SalusCall, while the probability of a miscalled base matching a random cluster is 25%. This indicates that spatial crosstalk accounts for a significant portion (∼25%) of the total miscalls. In contrast, AICall’s result shows that this percentage is only 25%-30%, indicating that the impact of spatial crosstalk has been reduced significantly. This benefits from our proposal to transform the base classification problem into an instance segmentation problem, where the entire fluorescence image is taken as input. This approach fully leverages the strengths of convolutional neural networks and GFSA in extracting spatial features, effectively eliminating the impact of spatial crosstalk on base classification. Meanwhile, since our method incorporates pixel information from a larger receptive field around the cluster, compared to other methods that only classify based on the center pixel of the cluster, this method also potentially mitigates classification errors caused by channel image registration misalignment.

### Phasing and pre-phasing

Phasing and pre-phasing occur when the incorporation of bases into the DNA strand is not synchronized (Fig. 4a). For all the clusters, if we plot their intensity at i+1 cycle vs. i cycle in a color channel, e.g. A channel, the scattered dots will again form 3 lines (Fig. 4b). The more horizontal line represents cycle i-A: cycle i+1-X case, which shows the typical phasing phenomenon where cycle i impacts cycle i+1’s call; the more vertical line represents cycle i-A: cycle i+1-A case, which shows the typical pre-phasing phenomenon where cycle i+1 impacts cycle i’s call; the diagonal represents both A case; and the origin spot represents neither A case.

**Fig. 4.**
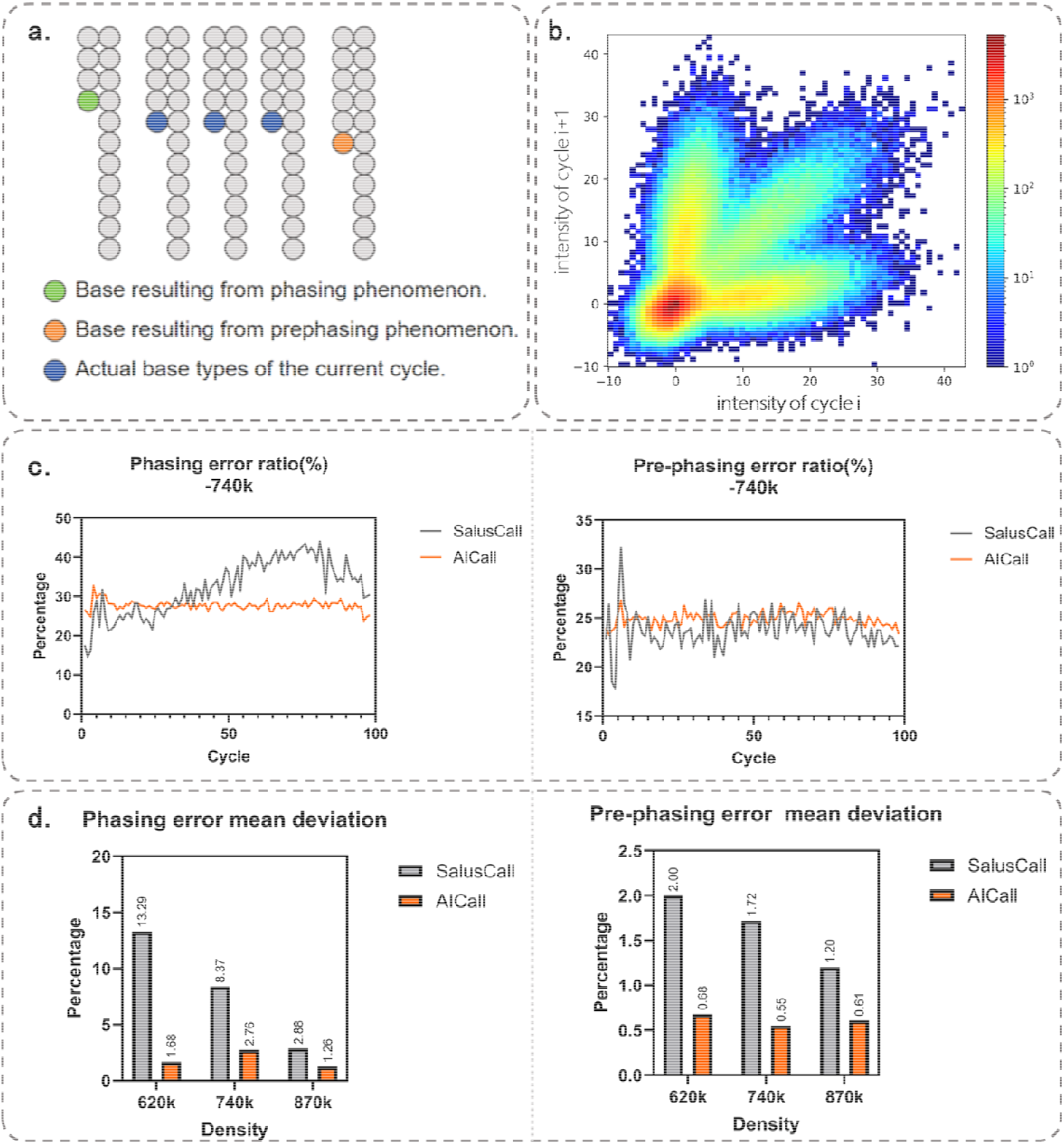
Phasing/pre-phasing correction. (a) Illustration of phasing and pre-phasing. (b) Statistics of cluster brightness in adjacent cycles for a certain color channel. (c) Error bias evaluated in terms of matching previous and subsequent base in each cycle for AICall and SalusCall. (d) Comparison of average deviation from the random probability (25%) for errors caused by phasing/pre-phasing using AICall and SalusCall.

Traditional basecall algorithm models the phasing/pre-phasing effect on the intensities of these cycles as straight lines. The “true” intensity of a cluster can then be obtained by removing the phasing/pre-phasing effects. However, the linear assumption may be an over-simplification. As a result, the correction is usually not satisfactory (Fig. 4c). For all miscalls from SalusCall, the proportion of those match previous ones becoming increasingly larger over sequencing cycles, from ∼20% to ∼40% (Fig. 4c left), with an average deviation of +8.37% from random probability (Fig. 4d left). This indicates that phasing problem is under-corrected and errors are more prone to happen at later cycles. As a contrast, AICall maintains a steady rate of ∼28% over all cycles, with an average deviation of +2.76%, showing its capability in capturing the characteristics of phasing problem. For pre-phasing problem, both SalusCall and AICall gave a percentage for those matching following base close to ∼25%, similar to the expected value (Fig. 3c right). The average deviation is only 0.2%∼2%. This indicates that pre-phasing is less of a problem as pre-phasing is usually associated with the purity of the chemical reagents (i.e., the dNTP losing the terminator group).

AICall demonstrates a clear theoretical advantage over traditional basecallers in addressing phasing and pre-phasing issues. Traditional basecallers typically estimate phasing/pre-phasing coefficients using least squares fitting and uniformly apply them to all clusters. In contrast, AICall classifies each cluster in the image by integrating pixel information from three consecutive cycles, enabling more precise modeling of the phasing/pre-phasing characteristics of the specific cluster. Furthermore, the GFCA module enhances feature representation by establishing inter-channel dependencies across cycles, while the SARB improves the model’s ability to learn the dynamic changes of phasing/pre-phasing effects over sequencing cycles. These advancements collectively enhance the robustness and accuracy of sequencing under the influence of phasing/pre-phasing interference.

### Spectral crosstalk and intensity-related phenomena

**Spectral crosstalk** is primarily caused by the emission spectrum overlap between fluorescent dyes marking different nucleotides. Fig.5a illustrates a case where an A cluster emission crossed into the T channel and showed as a bright spot in the T channel images. When we analyze all the clusters in two color channels e.g., T vs A channels, the scatter spots mainly form two lines. The more horizontal line represents the T clusters and the more vertical lin represents A clusters; the origin spot represents non-A-non-T clusters. As the crosstalk between the two color channels becomes more severe, the two lines will move toward each other.

Different instruments have their own signature of spectral crosstalk’s depending on the selection of fluorescent dyes and optic filters. Traditional algorithm corrects the spectral crosstalk from the measured crosstalk pattern. However, due to the random nature of errors, many clusters are distributed in the middle region of the crosstalk plot, making basecalls challenging for conventional basecallers such as SalusCall. Fig. 5c shows the A to T error ratio in all 12 types of single nucleotide substitution miscalls, where A and T spectral crosstalk is typically large for SalusPro instrument, with an average proportion reaching 22.7%. AICall consistently outperforms SalusCall, with an average error proportion of 13.5%, which is closer to the random expectation of 8.3%. AICall effectively mitigates the impact of spectral crosstalk on base classification, which is attributed to the GFCA module within its neural network. This module employs group convolutions and MLPs to capture the interdependencies among channel features within the same cycle, thereby optimizing the feature representations of different channels within the cycle. By enhancing the network’s robustness against spectral crosstalk, the GFCA module significantly improves the accuracy and reliability of base classification.

**Fig. 5.**
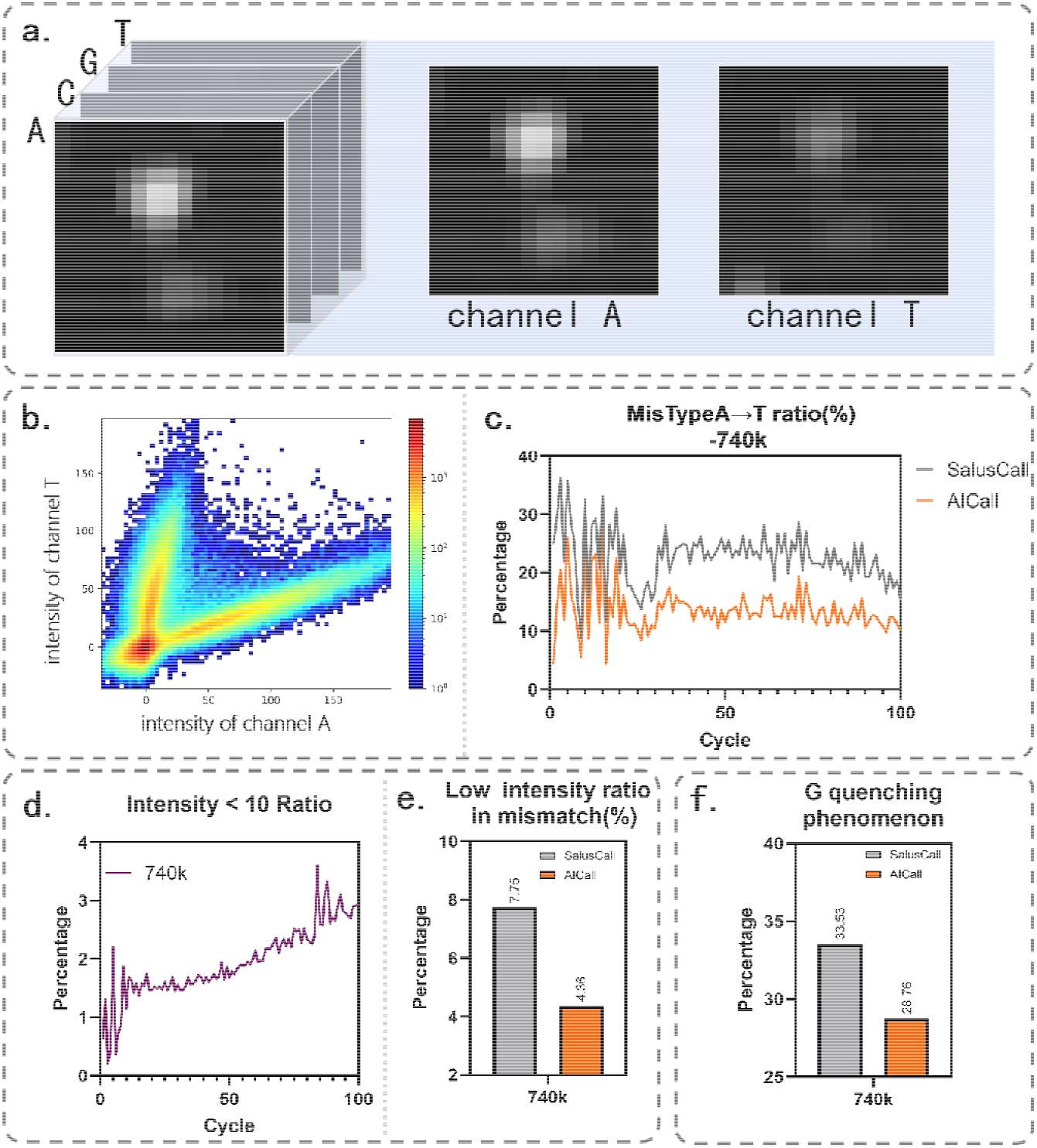
Spectral crosstalk and signal strength related basecalls. (a) Illustration of A to T crosstalk. (b) Scatter plots of clusters intensity in T channel vs in A channel. (c) The proportion of A miscalled as T by AICall and SalusCall. (d) The phenomenon of cluster intensity decay. (e) The proportion of low intensity clusters in mismatched clusters called by AICall and SalusCall. (f) The proportion of the previous base being G in the mismatch cluster called by AICall and SalusCall.

#### Fluorescent signal decay

As the number of sequencing reaction cycles increases, the signal quality of clusters deteriorates, which in turn affects basecall accuracy. <u>Fig. 5d shows that the proportion of low intensity clusters (intensity less than 10 ) increases as the cycle number increases</u>. We compared the mismatch rates of low-intensity clusters in AICall and SalusCall. As shown in Fig. 5e, the proportion of low-intensity clusters in the mismatch clusters called by AICall has significantly decreased. **G-quenching** occurs when the base in the previous cycle is G, leading to poorer signal intensity in the current cycle, which increases the likelihood of a misclassification. Again, Fig. 5f shows that, in the mismatch clusters of AICall sequencing results, the probability of the previous cycle being a G base is closer to the random probability of 25%, with a deviation of 3.78%. In contrast, SalusCall shows a probability of 33.53%, deviating by 8.53% from the random probability. AICall effectively reduces the bias towards low-intensity clusters in mismatches, which is attributed to the designed SEM and SARB. These modules encode cycle index information into high-dimensional vectors and embed them into the feature maps. By automatically adjusting the feature weights for each cycle, this mechanism enhances the neural network’s sensitivity to cycle-dependent variations in the images, such as fluorescence intensity decay and other related phenomena.

### Absolute error rate comparison

All above comparisons between methods are based on the basecall mismatch rate against the reference. However, since mismatch can be caused by de novo mutations during product generation of E.coli [10.1038/nature18313] and PCR errors in the library preparation etc., it does NOT reflect the error rate for the basecalls. To assess the absolute basecall errors, we adopted a mismatch against overlapped pairs (MAOP) method, in which errors are only called when the first and the second reads of pair-end sequencing are inconsistent .

We constructed a paired-end WGS sequencing library on E. coli (ATCC8739) and human genome (NA12878) using the Salus kit. Totally 297M sequenced bases for E. coli and 299M for human were approximately collected to calculate the mismatch rate. We conducted a comparison among the MAOP (Mismatch Against Overlapped Pairs) metric of AICall, SalusCall as well as Illumina. The sequencing results of AICall indicate that in both the E. coli and human samples, it has reached the lowest levels in terms of the MAOP metrics (Table 2). Notably, the AI call lowered the error rate to be less than 0.01%, ∼one order of magnitude better than current results. These findings indicate that AICall holds a substantial edge in sequencing precision, yielding more dependable and high-quality sequencing outcomes across various samples, which enhances the subsequent bioinformatics analysis.

**Table 2.** Results of the resampled Sequencing data from AICall, SalusCall and Illumina.

| Methods | MAOP error rate (%) |
| --- | --- |
| AICall | 0.0097 |
| SalusCall | 0.04 |
| Illumina | 0.04 |

## Discussion

This paper introduces AICall, a basecalling pipeline that integrates deep learning techniques and encompasses critical stages such as data preprocessing, training set construction, model design, and training. We proposed multiple metrics to analyze various interference factors in the sequencing process, including spatial crosstalk, spectral crosstalk, phasing, pre-phasing, intensity decay and G quenching. Extensive testing on datasets of varying densities has demonstrated that AICall significantly surpasses the traditional method SalusCall in sequencing accuracy. Notably, AICall exhibits enhanced robustness when confronting different interference factors, thereby providing more precise and reliable sequencing results for downstream analysis tasks.

AICall exhibits a marked advantage over traditional basecalling methods in managing complex interference factors in fluorescent images. The neural network model autonomously learns and extracts cluster features from large datasets, identifying more implicit patterns compared to manual feature design, thus offering enhanced robustness. When addressing dense clusters, the convolutional neural network constructs a broad receptive field through multiple downsampling and convolutional layers, allowing it to naturally account for the interference from multiple surrounding clusters, resulting in more accurate classifications. Moreover, AICall excels in handling channel-dimension interference. It effectively manages both intra-cycle spectral crosstalk and inter-cycle phasing and pre-phasing effects. During sequencing data processing, AICall initially converts the data into feature maps, with each feature map representing a channel. The model then uses encoders and decoders to expand and compress the channels, enabling the integration and adjustment of information across different channels. This process facilitates inter-channel communication and captures dependencies and correlations among them. By holistically considering information from multiple channels, the AICall model effectively reduces channel-dimension interference, ensuring accurate base classification.

The AICall model leverages deep learning techniques to automatically learn and extract useful features from raw sequencing data, significantly reducing the need for manual intervention and feature design. This innovation enables the AICall model to maintain strong adaptability across various platforms and different types of second-generation sequencers. As a result, the AICall model can quickly migrate to different sequencing platforms, using deep learning mechanisms to replace traditional sequencing methods and improve the accuracy and efficiency of data analysis. Thus, AICall’s exceptional performance not only highlights its robustness in managing complex data and interference factors but also demonstrates its broad applicability in diverse application scenarios.

AICall’s capability to accurately classify denser clusters enables the acquisition of more sequencing information within a smaller spatial area, which is significant for spatial genomics. The primary goal of spatial genomics is to obtain gene expression data from each position in a tissue slice using high-throughput sequencing technology, establishing a correspondence between gene expression and spatial location. Traditional low-resolution data may mix information from multiple cells, whereas high-density sequencing methods can enhance the resolution of spatial genomics, ensuring that each sequencing point primarily reflects information from a single cell, thus improving the accuracy of cell type identification. Additionally, higher sequencing resolution aids in more finely depicting tissue structure, such as distinguishing smaller cell populations and subcellular structures, revealing microscopic differences within tissues. High-density sequencing, therefore, provides robust technical support for spatial genomics research. Moving forward, we will further explore the relationship between AICall’s high-density sequencing and spatial genomics to propel the development of this field.

By integrating traditional methods with deep learning methods, AICall not only maximizes their respective advantages, enhancing processing speed, but also ensures prediction accuracy, providing an efficient and accurate solution for basecalling in fluorescent images.

## Acknowledgements

This work is partly funded by the Department of Natural Resources, China. Funding number: 2024ZD1000603.

## Reference

1. Cokus, S. J. et al. Shotgun bisulphite sequencing of the Arabidopsis genome reveals DNA methylation patterning. Nature 452, 215–219 (2008).

2. Pennisi, E. Breakthrough of the year. Human genetic variation. Sci. (New York, NY) 318, 1842–1843 (2007).

3. Begun, D. J. et al. Population genomics: whole-genome analysis of polymorphism and divergence in Drosophila simulans. PLoS Biol. 5, e310 (2007).

4. Nayanah, S. 1000 genomes project. Nat Biotech 26, 256 (2008).

5. Sanger, F., Nicklen, S. & Coulson, A. R. DNA sequencing with chain-terminating inhibitors. Proc. Natl. Acad. Sci. 74, 5463–5467 (1977).

6. Shendure, J. & Ji, H. Next-generation DNA sequencing. Nat. Biotechnol. 26, 1135–1145 (2008).

7. Pettersson, E., Lundeberg, J. & Ahmadian, A. Generations of sequencing technologies. Genomics 93, 105–111 (2009).

8. Chaisson, M. J. & Pevzner, P. A. Short read fragment assembly of bacterial genomes. Genome Res. 18, 324–330 (2008).

9. Bentley, D. R. et al. Accurate whole human genome sequencing using reversible terminator chemistry. Nature 456, 53–59 (2008).

10. Li, L. & Speed, T. P. An estimate of the crosstalk matrix in four-dye fluorescence-based DNA sequencing. Electrophoresis 20, 1433–1442 (1999).

11. Dohno, C. & Saito, I. Discrimination of Single-Nucleotide Alterations by G-Specific Fluorescence Quenching. ChemBioChem 6, 1075–1081 (2005).

12. Behlke, M. A., Huang, L., Bogh, L., Rose, S. D. & Devor, E. J. Fluorescence Quenching by Proximal G-bases. in (2005).

13. Erlich, Y., Mitra, P. P., delaBastide, M., McCombie, W. R. & Hannon, G. J. Alta-Cyclic: A self-optimizing base caller for next-generation sequencing. Nat. Methods 5, 679–682 (2008).

14. Rougemont, J. et al. Probabilistic base calling of Solexa sequencing data. BMC Bioinformatics 9, 1–12 (2008).

15. Wang, B., Wan, L., Wang, A. & Li, L. M. An adaptive decorrelation method removes Illumina DNA base-calling errors caused by crosstalk between adjacent clusters. Sci. Rep. 7, 1–11 (2017).

16. Zhou, Z., Siddiquee, M. M. R., Tajbakhsh, N. & Liang, J. UNet++: Redesigning Skip Connections to Exploit Multiscale Features in Image Segmentation. IEEE Trans. Med. Imaging 39, 1856–1867 (2020).

